# Predicting activatory and inhibitory drug–target interactions based on mol2vec and genetically perturbed transcriptomes

**DOI:** 10.1101/2021.03.18.436088

**Authors:** Won-Yung Lee, Choong-Yeol Lee, Chang-Eop Kim

## Abstract

A computational approach to identifying drug–target interactions (DTIs) is a credible strategy for accelerating drug development and understanding the mechanisms of action of small molecules. However, current methods are limited to providing simple DTIs without mode of action (MoA), or they show unsatisfactory performance for a limited range of compounds and targets. Here, we propose AI-DTI, a novel method that predicts activatory and inhibitory DTIs by combining the mol2vec and genetically perturbed transcriptomes. We trained the model on large-scale DTIs with MoA and found that our model outperformed a previous model that predicted activatory and inhibitory DTIs. To extend the applicability, we applied the inferential method for the target vector so that our method can learn and predict a wider range of targets. Our method achieved satisfactory performance in an independent dataset where the target was unseen in the training set and a high-throughput screening dataset where positive and negative samples were explicitly defined. Finally, our method successfully rediscovered approximately half of the DTIs for drugs used in the treatment of COVID-19. These results indicate that AI-DTI is a practically useful tool for guiding drug discovery processes and generating plausible hypotheses that can reveal unknown mechanisms of drug action.

## INTRODUCTION

Identifying drug–target interactions (DTIs) is an essential step in drug discovery and repurposing. Proper understanding of DTIs can lead to fast optimization of small molecules derived from phenotypic screening and elucidation of the mechanism of action for experimental drugs or natural products (1). However, identifying a candidate drug for a putative target by relying solely on *in vivo* and biochemical approaches often takes 2–3 years, with tremendous economic costs (2, 3). Computational prediction of DTIs has emerged as a promising new strategy for reducing the workload and resources needed by exploring enormous chemical and target spaces. This strategy has the potential to accelerate the drug development process by prioritizing candidate compounds for putative targets or *vice versa*.

Conventional methods for predicting DTIs can be broadly categorized into docking simulations and ligand-based approaches (4, 5). However, their prediction is often unreliable when the 3D structure of a protein or target is unavailable or when an insufficient number of ligands is known for the target (6). Recently, chemical genomic approaches have emerged as an alternative enabling large-scale predictions by leveraging recent advances in network-based approaches or machine learning techniques (7–11). For example, Yunan et al. proposed DTI-Net, a network-integrated pipeline that predicts DTIs by constructing a heterogeneous network using the information collected from various sources (12). Other researchers have proposed deep learning-based methods, such as CNN, GCN, and NLP, to predict novel DTIs (13–15). Despite their state-of-the-art performance, these models predict simple interactions without the mode of action, providing limited guidance for drug development or necessitating further experimental validation to fully understand the mechanisms of action of the drug.

Genetically perturbed transcriptomes are a valuable resource that can comprehensively measure the response of a biological system following genetic perturbations. Recently, the LINCS Program launched L1000, a new, low-cost, high-throughput, reduced representation expression profiling method with a collection of more than one million gene expression profiles (16, 17). Using such a large-scale dataset, Sawada et al. developed a model that predicts activatory and inhibitory DTIs by combining transcriptome profiles measured after compound treatment and genetic perturbation (18). Although the method showed the possibility of predicting DTIs with modes of action using transcriptome data, it did not provide a satisfactory tool that could be applied for drug discovery. First, employing a compound vector to represent the transcriptome profiles measured after compound treatment limited the range of predictable compounds significantly, making it difficult to apply the model to predict DTIs for experimental drugs and natural products. Second, the number of predictable activatory and inhibitory targets was 74 and 755, respectively, which covered only a fraction of the druggable targets. Finally, the employed algorithms, called joint learning, could not learn nonlinear relationships between input vectors and DTI labels, and thus, the performance of DTI prediction was insufficient. Therefore, there is still a pressing need to develop a method with superior performance for predicting a wide range of activatory and inhibitory targets for novel compounds and natural products.

In this paper, we present AI-DTI, a new computational methodology for predicting activatory and inhibitory DTIs, by integrating a mol2vec method and genetically perturbed transcriptomes (Figure 1). Employing mol2vec enabled our method to expand the drug space to any compound for which extended connectivity fingerprints (ECFPs) can be calculated. In addition, the target space of our method was expanded to cover a majority of druggable targets by inferring the target vector representation based on the protein–protein interaction (PPI) network. AI-DTI achieved substantial performance improvement over the previous model in both activatory and inhibitory DTI predictions. The performance was still satisfactory for independent datasets where protein target was unseen in the training set and where positive and negative samples were explicitly defined. Finally, as a case study, we confirmed the successful rediscovery of more than half of the recently validated DTIs for COVID-19 via AI-DTI. All these results demonstrate that AI-DTI is a practically useful tool for predicting unknown activatory and inhibitory DTIs, which may provide new insights into drug discovery and help in understanding modes of drug action.

**Figure 1.**
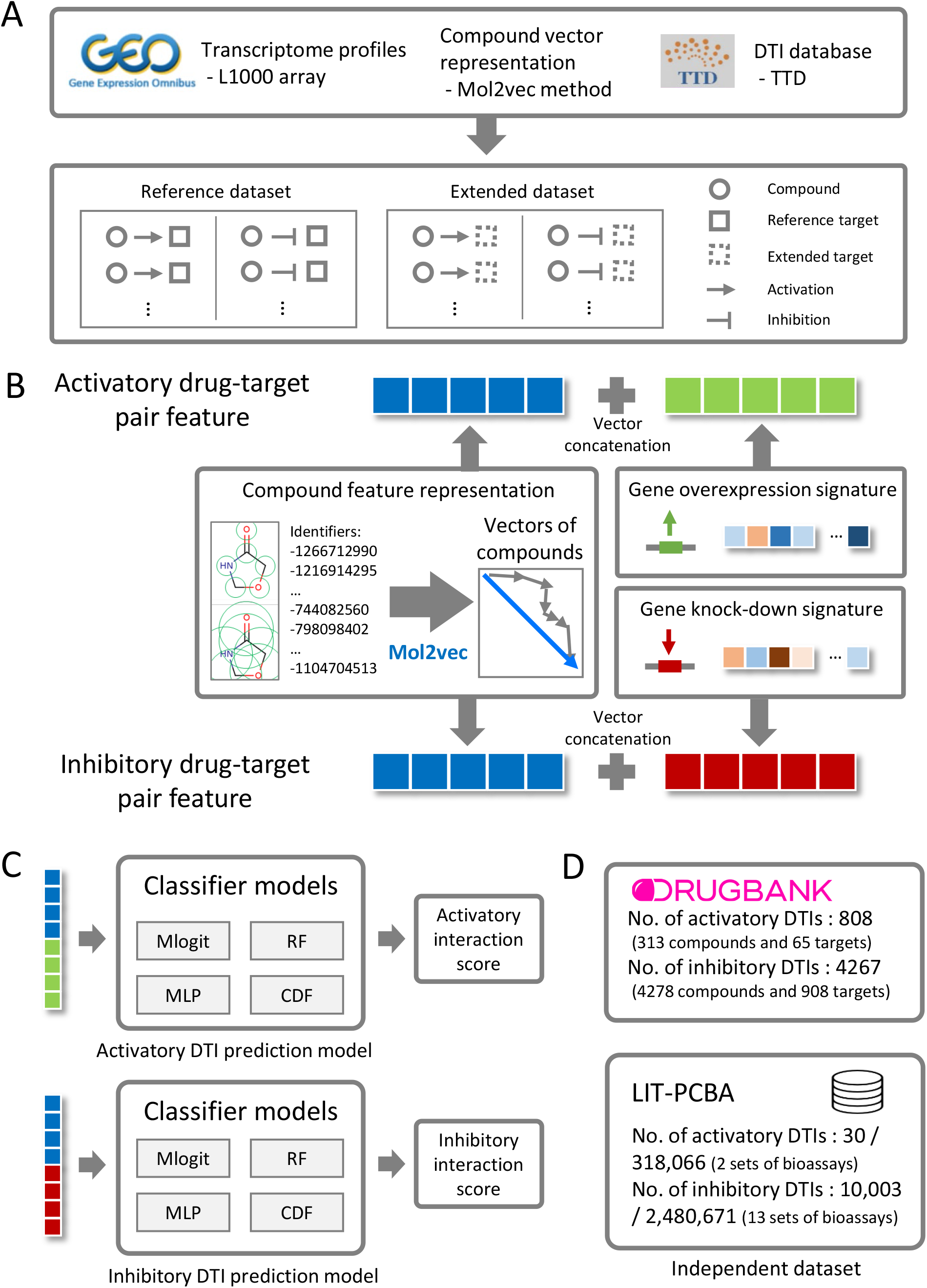
Overview of the AI-DTI pipeline. (A) Construction of a dataset of known DTIs. Transcriptome profiles were obtained from the L1000 array data and then aggregated to generate a representative target vector. A mol2vec method was used to generate representative vectors for compounds. DTIs with modes of action were collected from the TTD. The reference dataset was constructed by selecting activatory and inhibitory DTI pairs that include a compound for which ECFPs can be calculated and a reference target (i.e., a target for which genetically perturbed transcriptome data are available). The expanded dataset was constructed by selecting activatory and inhibitory DTI pairs that include a compound for which ECFPs can be calculated and an extended target (i.e., a target for which inferred transcriptome data are available). (B) Feature vector generation for activatory and inhibitory DTIs. For activatory DTIs, the feature vector was represented as a concatenation of the compound vector and aggregated gene overexpression signatures. For inhibitory DTIs, the feature vector was represented as a concatenation of the compound vector and aggregated gene knockdown signatures. (C) Classifier training process. The feature vector corresponding to the DTIs was fed into the classifier models to predict novel activatory and inhibitory DTIs. (D) External validation process. The fidelity of classification was evaluated in two independent datasets, viz., DrugBank and LIT-PCBA.

## MATERIAL AND METHODS

### Employing mol2vec-based compound features

The vector representation of compounds was obtained using mol2vec (19), a word2vec-inspired model that learns the vector representations of molecular substructures. Mol2vec applies the word2vec algorithm to the corpus of compounds by considering compound substructures derived from the Morgan fingerprint as “words” and compounds as “sentences”. The vector representations of molecular substructures are encoded to point in directions similar to those that are chemically related, and entire compound representations are obtained by summing the vectors of the individual substructures. Among the mol2vec versions, we implemented the skip-gram model with a window size of 10 and 300-dimensional embeddings of Morgan substructures, which demonstrated the best prediction capabilities in several compound property and bioactivity datasets.

### Constructing genetically perturbed transcriptome-based target features

The genetically perturbed transcriptome of the L1000 dataset was downloaded from the Gene Expression Omnibus (accession number: GSE92742), which contains 473,647 signatures. Each signature (i.e., transcriptome profile) consists of a mediated z-score of 978 landmark genes whose expression levels were measured directly and 11,350 genes whose expression values were inferred from them. Landmark gene refers to one whose gene expression has been determined as being informative to characterize the transcriptome and which is measured directly in the L1000 assay. In our study, level 5 landmark gene data were used to represent the target vector. Level 5 data are a normalized dataset suggested by the LINCS team for use without additional processing. Among the types of perturbations, “cDNA for overexpression of wild-type gene” and “consensus signature from shRNAs for loss of function” were considered vector representations for activatory and inhibitory targets, respectively. From the downloaded data, the gene expression signatures of the landmark gene set were parsed using the cmapPy module (20), resulting in 36,720 gene knockdown and 22,205 gene overexpression signatures.

The parsed data contained multiple gene expression profiles for single genetic perturbations measured in various cell lines and/or under perturbing conditions, which necessitated further preprocessing. To generate representative target vector by each target, we applied the weighted average procedure, a method for aggregating multiple signatures with weights based on a pairwise correlation matrix (Figure 2A) (16, 21). Suppose that **X**_t_∈ ℝ^978^ (t = 1, 2…, n) is a vector representing each L1000 signature of a landmark gene (normalized gene expression profiles for directly measured genes) for a specific functional perturbation, where t indicates a specific combination of cell lines and experimental conditions, and n indicates the total number of combinations. To generate a single representative vector **X**_*Rep*_, a pairwise correlation matrix **R**^**n**×**n**^ is defined as the Spearman coefficient between signature pairs,

**Figure 2.**
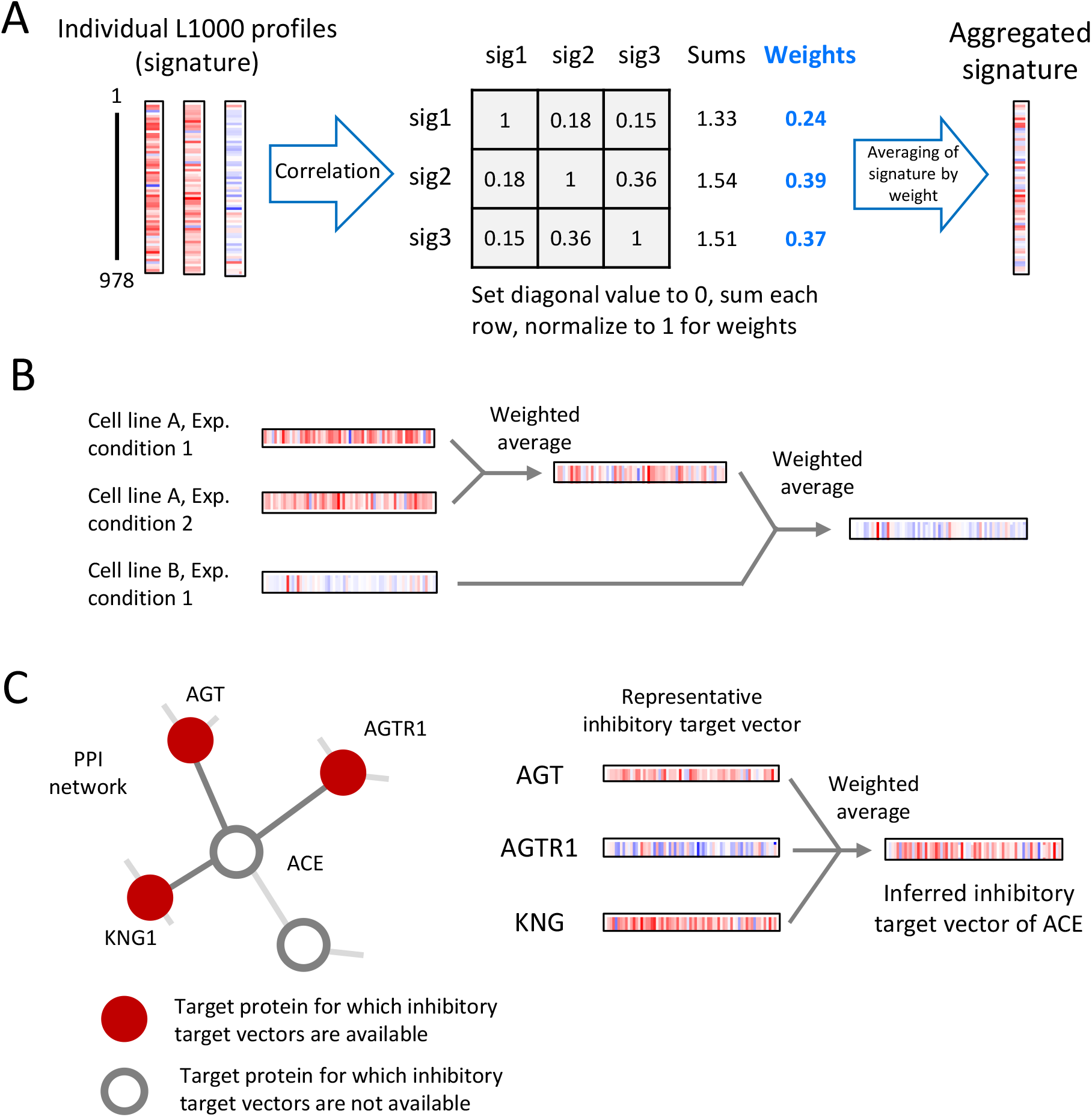
Schematics of aggregation and inference for a genetically perturbed transcriptome. (A) Weighted averaging for combining individual signatures into consensus gene signatures. Individual profiles were weighted by the sum of their correlations to other individual signatures and then averaged. (B) Generation of target vectors by cross-perturbation and cross-cell line aggregation. Multiple signatures measured after the same genetic perturbation in a specific cell, but with different perturbational doses or times, were first aggregated by weighted averaging. The same procedure was performed between multiple signatures measured after the same genetic perturbation in different cell lines. (C) Examples of inferring target vector representation. AGT, AGTR1, and KNG1 interacted with ACE in the PPI network, and their inhibitory target vectors were available. The inhibitory target vector of ACE was inferred by aggregating these three representative inhibitory target vectors using weighted averaging.

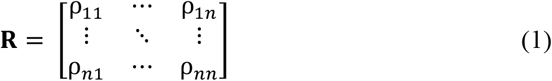

where ρ_*ij*_denotes the Spearman correlation coefficient between the signature pairs **X**_i_and **X**_j_.

The weight vector (**w**) is obtained by summing across the columns of **R** after excluding trivial self-correlation and then normalizing them,

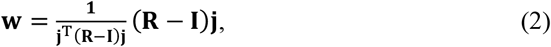

where **I** denotes the identity matrix, and **j** denotes column vectors of 1s. Finally, **X**_*Rep*_ is obtained from the average of **X**_*t*_ based on the weight vector **w**,

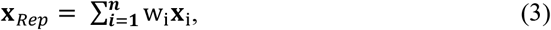

where w_i_ denotes the i-th entries of **w**.

The process of generating the representative target vector by applying the weighted average was divided into two step: aggregation across experimental conditions and aggregation across cell lines (Figure 2B). Specifically, the signatures measured after the same genetic perturbation in a specific cell, but with different perturbational dose or time, were first aggregated by weighted averaging. The representative vector for a particular target is obtained by reapplying weighted averaging to these signatures (i.e., the aggregated signatures measured after the same genetic perturbation in different cell lines). This segmentation process reduces the potential biases on the representative vector calculation that occurs when the number of genetically perturbed signatures skewed on a particular cell line. As a result, we obtained the representative vectors of 3,114 and 4,345 activatory and inhibitory targets, respectively, encoded in 978 dimensions (Figure 2B). The obtained target vector representation was used as target features of DTIs in a *reference dataset* to be constructed later.

The target list of the obtained vector contained only a fraction of the druggable targets, thus significantly limiting the target space of our method. Therefore, the target space was extended by inferring the vector representation of activatory or inhibitory signatures based on the PPI network. The PPI network was constructed from the STRING database (v 11.0) (22) by setting the organism as “homo sapiens” and an interaction score > 0.9 (highest confidence score suggested by STRING). The vector representation of the activation or inhibition target was inferred by aggregating the vector representation of the interacting protein in the PPI network using a weight averaging procedure (Figure 2C). To ensure the quality of the data, we limited the inferred targets to proteins with at least three neighbours whose target vectors are available. The inferred target vector representation was used as target features of DTIs in an *extended dataset* to be constructed later.

### Selecting inhibitory and activatory DTIs

Known activatory and inhibitory DTIs were obtained from the Therapeutic Target Database (TTD) 2.0 (accessed October 15, 2020) (23) and DrugBank 5.1.7 (accessed January 12, 2021) (24). We selected DTIs that explicitly defined activatory or inhibitory interactions (“activator” or “agonist” for activatory DTIs and “inhibitor” or “antagonist” for inhibitory DTIs). Identifiers of compounds and targets with their annotations were standardized by PubChem CID and gene symbols, respectively. The chemical structures of our dataset were retrieved in canonical SMILE format using the Python package PubChemPy. For TTD, we obtained 2,925 activatory and 32,417 inhibitory DTIs between 24,145 compounds and 2,117 targets. For DrugBank, we obtained 919 activatory and 4,719 inhibitory DTIs between 1,600 compounds and 1,022 targets.

### Training classifier models

The features of activatory and inhibitory DTIs were fed into classifier models to predict their interactions. The DTI prediction performance was evaluated by multinomial logistic regression (Mlogit), random forest (RF), multilayer perceptron (MLP), and cascade deep forest (CDF). RFs are ensemble models that combine the probabilistic predictions of a number of decision tree-based classifiers to improve the generalization capability over a single estimator. MLP is a supervised learning algorithm that can learn nonlinear models. The architecture and hyperparameters of the MLP models in this study are summarized in Supplementary Figure 1. CDF employs a cascade structure, where the model consists of a multilayered architecture, and each level consists of an RF and extra trees (25). Each level of cascade receives feature information processed by the preceding level and conveys its result to the next level. A key feature of the updated CDF model, deep forest 21, is that it automatically determines the number of cascade levels that are suitable for the training data by terminating the training procedure when performance improvements through adding cascade levels are no longer significant (26). Greedy-search methods were used to select optimized hyperparameters of CDF as follows: the number of estimators in each cascade layer, the number of trees in the forest, the function to measure the quality of a split, maximum depth, minimum number of samples to be split, maximum features, and minimum impurity decrease. The detailed search range of the optimization is summarized in Supplementary Table S1.

### Performance evaluation

The performance was evaluated using fivefold cross-validation (CV). For each fold of the predictive model, the following metrics were calculated:

Precision = TP/TP + FP

True positive rate = TP/TP + FN

False positive rate = FP/FP + TN,

where TP is true positive, FP is false positive, FN is false negative, and TN is true negative. We plotted the receiver operating characteristic (ROC) curves based on different recall and false positive rates and precision-recall (PR) curves based on different precision and recall values under the conditions of different classification cut-off values. The area under the ROC curve (AUROC) and area under the PR curve (AUPR) were calculated over each fold, and their average values were recorded as measures of model performance. AUPR provides a better assessment in highly skewed datasets, whereas AUROC is prone to be an overoptimistic metric. Thus, we used AUPR as the key metric for model selection (27, 28). To evaluate the performance in the specific threshold in the ROC curve, the enrichment in true actives at a constant x% false positive rate over random picking (EFx%) was calculated as follows:

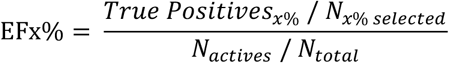

The EFx% value represents the enriched true positive rate compared to the expected value at the threshold, representing a false positive rate of x%. For example, the EF1% value indicates the enrichment of the true positive rate compared to the chance level (0.01) at the threshold where the false positive rate is 0.01.

## RESULTS

### Overview of AI-DTI

We developed AI-DTI for the *in silico* identification of activatory and inhibitory DTIs (Figure 1). To conduct supervised learning, our method constructs the input features of DTIs by concatenating the vectors of compounds and targets derived from the mol2vec method and genetically perturbed transcriptome, respectively. The mol2vec method transforms the structural information of a 2D compound into a continuous multidimensional vector, and a genetically perturbed transcriptome embeds the response of the biological system following gene overexpression or silencing, which is associated with the response when drugs activate or inhibit specific gene targets (29–31). We constructed the input features for predicting activatory and inhibitory DTIs, respectively. For activatory DTIs, the feature vector was represented as a concatenated form of the compound vector calculated by mol2vec and the representative vectors of activatory targets. For inhibitory DTIs, the feature vector was represented as a concatenated form of the compound vector calculated by mol2vec and the representative vectors of inhibitory targets (see *Materials and Methods*). Based on the constructed dataset consisting of the DTIs and its input features, we trained classifier models and selected the model with the best performance to predict activatory and inhibitory DTIs. The generalized ability of the selected model was measured using two independent datasets, viz., DrugBank and LIT-PCBA (32).

### AI-DTI accurately predicts activatory and inhibitory DTIs

We first constructed a dataset, referred to as the “Reference dataset”, by selecting a known pair of activatory and inhibitory DTI pairs that include a compound for which ECFPs can be calculated and a target for which transcriptome data are available (Table 1). We trained the Mlogit, RF, MLP, and CDF models on the reference dataset and then evaluated the performance by fivefold CV. In each fold, a training set was constructed by randomly selecting a subset of the 80% known DTI pairs (assigned as the positive sample) and the matching number of randomly selected DTI pairs (assigned as the negative sample), and the test set was constructed by selecting the remaining 20% of the known DTI pairs and a matching number of randomly selected DTI pairs. To reduce the data bias of cross-validation, we performed 5 random 5-fold CVs and evaluated the performance of the model. Our results showed that the CDF model yielded the highest AUROC and AUPR values in both situations when predicting activatory or inhibitory targets (Table 2). We subsequently tried to optimize the CDF model and found that the highest AUROC and AUPR values were obtained in all situations when the following hyperparameters were selected: ‘500’ as the number of trees and ‘8’ as the number of estimators (Supplementary Table 1). The optimized CDF model achieved AUROC and AUPR values of 0.880 and 0.899 for predicting activatory DTIs and 0.935 and 0.946 for predicting inhibitory DTIs, respectively (Table 3).

**Table 1.** Overview of the drug–target interaction dataset for model training and external validation.

|  | Activatory DTIs |  |  | Inhibitory DTIs |  |  |
| --- | --- | --- | --- | --- | --- | --- |
|  | No. of compounds | No. of targets | No. of DTIs | No. of compounds | No. of targets | No. of DTIs |
| Internal sets |  |  |  |  |  |  |
| Reference dataset | 457 | 87 | 513 | 6,789 | 702 | 8,328 |
| Extended dataset | 887 | 186 | 1,242 | 6,781 | 332 | 9,545 |
| Total | 1,265 | 273 | 1,755 | 12,259 | 1,034 | 17,873 |
| External sets |  |  |  |  |  |  |
| DrugBank* | 374 | 172 | 808 | 1,217 | 909 | 4,267 |
| LIT-PCBA | 130,412 | 2 | 30/318,066<sup>#</sup> | 302,567 | 13 | 10,003/2,480,671<sup>#</sup> |

\*A dataset with only DTIs unseen in the TTD dataset. <sup>#</sup>Number of active and inactive DTIs.

**Table 2.**
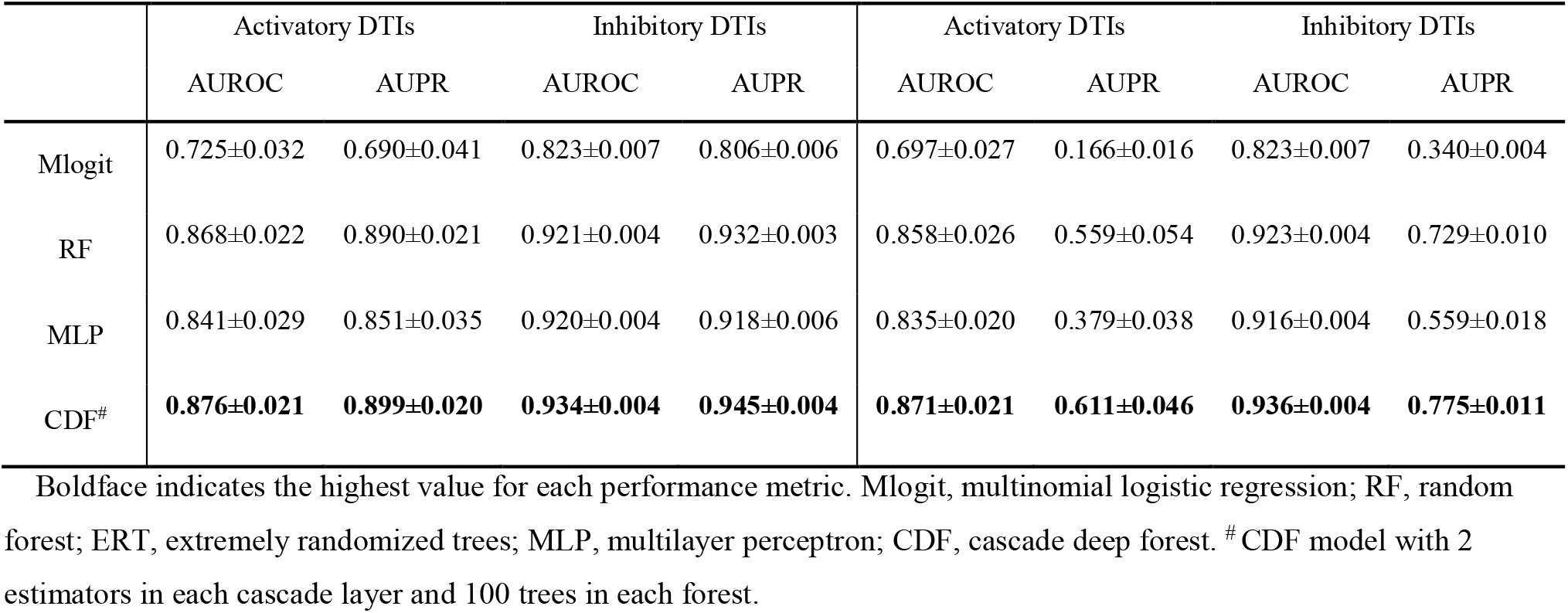
Assessment of performance using the reference datasets through fivefold cross-validation.

| Sample ratio = 1:1 (mean±S.D) |  |  |  |  | Sample ratio = 1:10 (mean±S.D) |  |  |  |
| --- | --- | --- | --- | --- | --- | --- | --- | --- |
|  | Activatory DTIs |  | Inhibitory DTIs |  | Activatory DTIs |  | Inhibitory DTIs |  |
|  | AUROC | AUPR | AUROC | AUPR | AUROC | AUPR | AUROC | AUPR |
| Mlogit | 0.725±0.032 | 0.690±0.041 | 0.823±0.007 | 0.806±0.006 | 0.697±0.027 | 0.166±0.016 | 0.823±0.007 | 0.340±0.004 |
| RF | 0.868±0.022 | 0.890±0.021 | 0.921±0.004 | 0.932±0.003 | 0.858±0.026 | 0.559±0.054 | 0.923±0.004 | 0.729±0.010 |
| MLP | 0.841±0.029 | 0.851±0.035 | 0.920±0.004 | 0.918±0.006 | 0.835±0.020 | 0.379±0.038 | 0.916±0.004 | 0.559±0.018 |
| CDF<sup>#</sup> | **0.876±0.021** | **0.899±0.020** | **0.934±0.004** | **0.945±0.004** | **0.871±0.021** | **0.611±0.046** | **0.936±0.004** | **0.775±0.011** |

Boldface indicates the highest value for each performance metric. Mlogit, multinomial logistic regression; RF, random forest; ERT, extremely randomized trees; MLP, multilayer perceptron; CDF, cascade deep forest. <sup>#</sup>CDF model with 2 estimators in each cascade layer and 100 trees in each forest.

**Table 3.** Assessment of the performance of the optimized CDF model for various datasets.

| Sample ratio = 1:1 (mean±S.D) |  |  |  |  | Sample ratio = 1:10 (mean±S.D) |  |  |  |
| --- | --- | --- | --- | --- | --- | --- | --- | --- |
|  | Activatory DTIs |  | Inhibitory DTIs |  | Activatory DTIs |  | Inhibitory DTIs |  |
|  | AUROC | AUPR | AUROC | AUPR | AUROC | AUPR | AUROC | AUPR |
| Reference dataset | 0.880±0.029 | 0.899±0.019 | 0.935±0.003 | 0.946±0.003 | 0.873±0.007 | 0.629±0.033 | 0.939±0.004 | 0.780±0.007 |
| Extended dataset | 0.873±0.011 | 0.869±0.013 | 0.953±0.002 | 0.957±0.002 | 0.864±0.012 | 0.430±0.030 | 0.955±0.002 | 0.800±0.008 |
| Separate model* | **0.875±0.011** | 0.878±0.011 | **0.944±0.002** | **0.952±0.002** | 0.867±0.009 | 0.488±0.022 | **0.947±0.002** | **0.790±0.004** |
| Integrated model<sup>#</sup> | **0.875±0.010** | **0.881±0.008** | 0.943±0.002 | 0.951±0.001 | **0.869±0.008** | **0.489±0.023** | 0.946±0.002 | 0.786±0.005 |

Boldface indicates the highest value for each performance metric between the separate model and integrative model.
\*Models trained on the reference and extended datasets separately. <sup>#</sup>Models trained on the integrated dataset comprising the reference and extended datasets.

DTI prediction is an unbalanced classification problem where positive labels are sparse, so the performance measured on the dataset in which positive and negative samples are balanced does not fully reflect the situations in real drug discovery scenarios. To mimic the practical situation in which positive DTI is sparse, we also performed an additional cross-validation test, in which the negative set in the test data contained ten times more negative samples than positive samples. With this experimental setup, the known DTI (i.e., positive samples) accounts for only 9% of the total data set, allowing a performance assessment closer to the situation of real drug discovery. Although the scores dropped when compared to the previous test, we observed that the optimized CDF and RF models still achieved high AUPR values (Table 2). The AUPR of the MLP model was significantly lower than that of the above two models, indicating that the performance of the MLP model was insufficient in the skewed dataset. Considering the highest performances in the experimental setup, we decided to employ the optimized CDF model as the classifier model of AI-DTI.

To clarify the superiority of the approach, we compared the performance of the AI-DTI with the previous model for predicting activatory and inhibitory DTIs (18). The performance was evaluated under the same conditions as the previous work, which involved performing fivefold CV by assigning a negative dataset with all remaining nonpositive DTIs. We trained and predicted DTIs using the optimized CDF model that showed the best performance in the above experiments and Mlogit with l1 regularization, the baseline model employed in the previous study. We observed that our method still had much higher performances than previous models in both situations for predicting activatory and inhibitory DTIs (Table 4). The improved performance of the CDF model over the previous method indicates that our approach learned the relationships of activatory and inhibitory interactions between compounds and targets more efficiently than joint learning. Furthermore, we observe that our method using the Mlogit model with l1 regularization achieves better performance in terms of AUROC than the previous model based on the same model, indicating that the feature representation of our method provides a more accurate granular task classification ability. We observed that Mlogit method showed lower AUPR compared to the previous model, but this is because the positive samples in our dataset are sparser than the previous study, which is a more unfavorable condition for the AUPR metric.

**Table 4.**
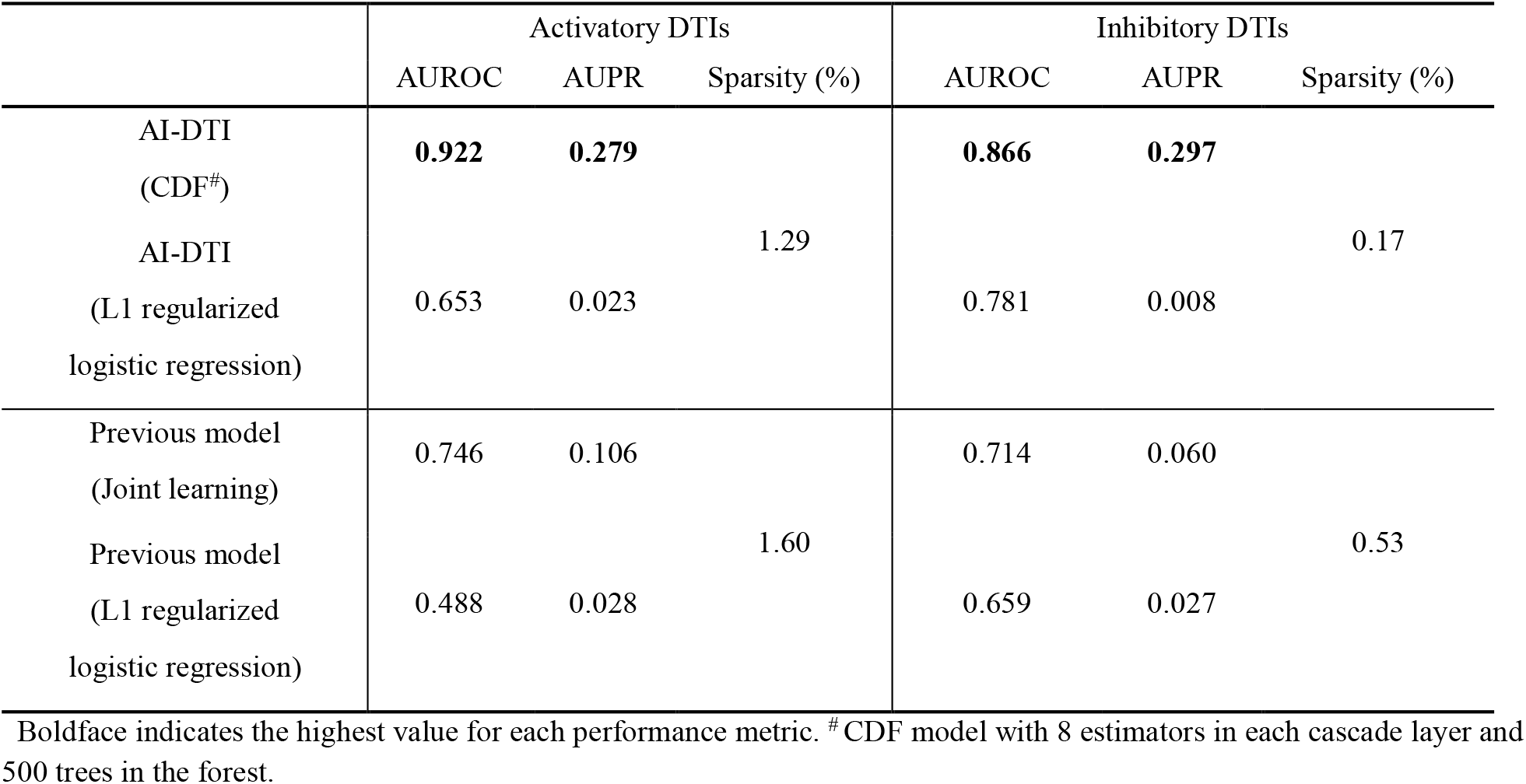
Comparison of performance for our model and previous models under the same experimental procedure.

### AI-DTI can predict diverse druggable targets

The drawback of previous models using genetically perturbed transcriptomes is that the range of predictable targets is constrained to the targets for which the genetically perturbed transcriptome is measured. For example, in a previous study (18), the number of predictable activatory and inhibitory DTI targets was only 77 and 769, respectively, covering only a fraction of druggable targets. To broaden the applicability of our method, we attempted to expand the target space of our model by inferring target vectors based on PPI networks (see *Materials and Methods*, Figure 1C). The assumption for using this method is that genetically perturbed transcriptomes are correlated with those of functionally interacting proteins. The inferring procedure calculates a representative vector for a target whose genetically perturbed transcriptome was not measured, thus enabling the construction of the input feature of DTI pairs for wider targets. We constructed another dataset, referred to as the “Extended dataset”, by selecting DTI pairs that include a compound for which ECFPs can be calculated and a target for which inferred transcriptome data are available (Table 1). The extended dataset secured an additional 186 and 332 targets for activatory and inhibitory DTIs, respectively, resulting in our dataset covering more than 70% of druggable targets appearing in the TTD. The performance of the optimized CDF model was used to evaluate the fidelity of the embeddings of DTIs in the extended dataset. The results showed that the CDF model achieved satisfactory AUROC and AUPR values in the extended dataset (Table 3), indicating that activatory and inhibitory DTIs can be accurately predicted even with the inferred target vectors. We observed that there was no significant change in the performance when training the model by integrating the reference dataset and extended dataset and when training the model using a separate dataset (Table 3). For ease of use, we decided to train the model using the integrated dataset and conducted subsequent analyses.

### AI-DTI performed satisfactorily for the independent dataset

The performance of AI-DTI was further evaluated on independent datasets. We obtained DTIs along with the mode of action from DrugBank and selected a DTI that meets the following two criteria: (1) DTI pairs that include a compound for which ECFPs can be calculated and a target for which (inferred) transcriptome profiles are available and (2) DTIs that were not seen during the training phase (Table 1). All the remaining nonpositive samples between drugs and targets were assigned to the negative samples. We found that the targets of DTIs in the dataset were completely unseen during the training phase, indicating that the constructed dataset can evaluate the generalizability of AI-DTI to identify DTI pairs in which the target was unseen in the training phase. We predicted activatory and inhibitory DTIs using the optimized CDF model trained on the integrated dataset. The results showed that our models achieved satisfactory AUROC and AUPR values, indicating the generalizability of our model to predict DTIs in which the targets were unseen during the training phase. Specifically, the optimized CDF-based model achieved the highest AUROC of 0.773 for activatory DTIs and 0.723 for inhibitory DTIs. The precision-recall curves revealed that the performance of the optimized CDF model was still better than that of the other models (Figure 3A).

**Figure 3.**
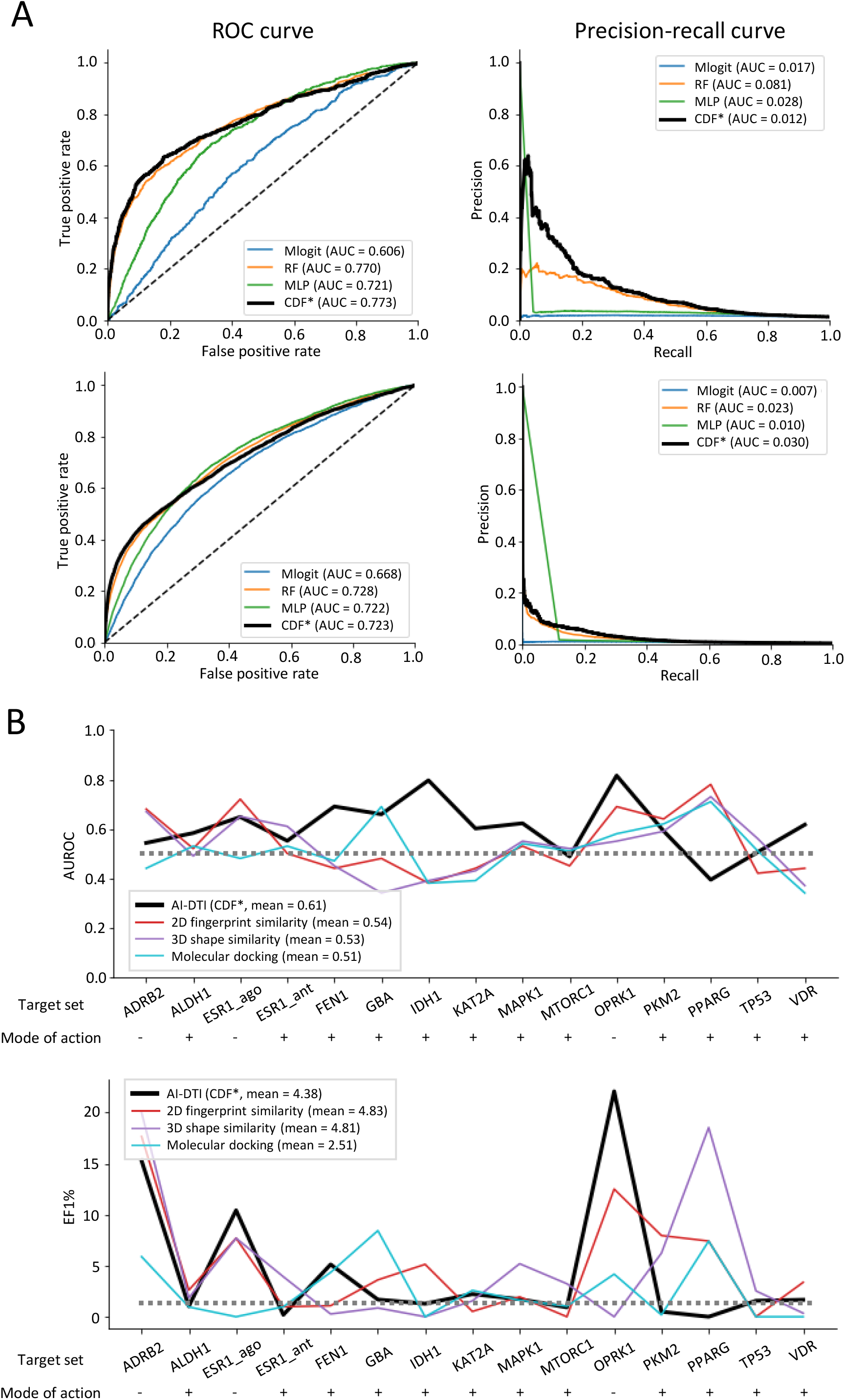
Assessment of performance on independent datasets. (A) Performance curves for activatory (top) and inhibitory (bottom) DTIs on the DrugBank dataset. ^*^CDF model with 8 estimators in each cascade layer and 500 trees in each forest. Mlogit, multinomial logistic regression; RF, random forest; MLP, multilayer perceptron. CDF, cascade deep forest. (B) Performance comparison between our optimized model and the conventional virtual screening methods on LIT-PCBA. ^*^ CDF model with 8 estimators in each cascade layer and 500 trees in each forest.

Evaluating the performance on the dataset by assigning nonpositive samples as negative samples does not fully reflect practical drug discovery scenarios. To this end, we evaluated the performance of AI-DTI in another benchmark dataset, LIT-PCBA, where negative samples and interaction types are explicitly defined (32). LIT-PCBA provides a comprehensive list of true active and true inactive compounds on high-throughput screening assay results for 15 targets (3 activatory targets and 12 inhibitory targets) (Table 1). We confirmed that all DTI pairs except for one active compound (bisindolylmaleimide i for the MAPK1 inhibitor) and all DTI pairs for the four target sets (ALDH1, FEN1, KAT2A and PKM2) in LIT-PCBA were not completely exposed during the training phase. These independencies from the training dataset indicate that this dataset can evaluate the generalized availability of AI-DTI for high-throughput screening datasets with unseen targets and/or compounds. We predicted activatory and inhibitory DTIs using the optimized CDF model trained on the integrated dataset and compared the performance with the baseline three virtual screening (VS) methods presented by LIT-PCBA, i.e., the 2D fingerprint similarity method, 3D shape similarity method, and molecular docking. To reduce the bias of the performance, we trained our methods ten times using different negative samples and measured the mean performance on a fully processed target set. The results show that AI-DTI achieved a higher mean AUROC than those achieved by conventional VS methods optimized with a max-pooling approach (Figure 3B). Specifically, our method achieved the highest AUROC values in all target sets (FEN1, IDH1, KAT2A, and VDR) where the other VS methods produced worse AUROC values than chance and for three target sets (ALDH1, FEN1, and KAT2A) that were unseen in the training phase. Moreover, our method achieved higher EF1% values than conventional methods for all activatory ligands (ESR_ago, FEN1, and OPRK1) and one inhibitory ligand (FEN1). It is worth noting that AI-DTI is a large-scale method that can predict a wide range of targets, whereas the other comparative models are local models built separately to predict specific protein targets. In summary, we found that our model still offers superior performance for classifying active and inactive compounds in high-throughput screening datasets containing DTIs with an unseen target and/or an unseen compound.

### AI-DTI can predict DTIs for novel diseases

To test the applicability for novel diseases, we tested whether AI-DTI could identify the DTIs of candidate drugs for COVID-19 treatment. Validated DTIs for COVID-19 were collected from DrugBank, and DTIs that met two criteria were further selected as follows: (1) DTIs containing a compound for which ECFPs could be calculated and a target with (inferred) transcriptome profiles available and (2) DTIs that were not seen during the training phase. We employed optimized CDF-based models trained on integrated datasets to predict the activatory or inhibitory interaction scores for validated DTIs. We regard the validated DTI to be rediscovered when the predicted score exceeds the default threshold of our model (0.5). To assess how uncommon the predicted score is, we constructed a reference distribution and compared a relative rank of the score (i.e., top %) to the reference distribution of interaction scores. The reference distribution was defined as the distribution of interaction scores calculated using our method for DTI pairs between 2500 FDA-approved drugs and a target of interest.

We found that approximately half of the DTIs (12/25) were successfully rediscovered by our method, of which three and five activatory and inhibitory DTIs were found to be in the top 5%, respectively (Table 5). It is noteworthy that the targets of two activatory DTIs (metenkephalin - OPRM1 and metenkephalin - OPRM1) and two inhibitory DTIs (ifenprodil - GRIN1 and ifenprodil - GRIN2B) were included in the extended dataset, so the high top percentages of these DTIs support the reliability of our results within the extended target space. On the other hand, the low true positive rate of the inhibitory DTIs (33%, 6/18) might raise concerns about the reliability of our method’s prediction results. However, except for the three targets (TNF, HMGB1m and JAK1), we found that all DTI scores between the FDA-approved drugs and targets showing false-negative results did not exceed the default threshold, which indicates that these false-negative results did not affect the precision of the predicted results. To facilitate drug repurposing, we used our method to summarize a list of FDA-approved drugs yielding high scores for COVID-19 targets (Supplementary Table S2-3). Taken together, these results demonstrate that AI-DTI can be used to identify DTIs in novel diseases.

**Table 5.**
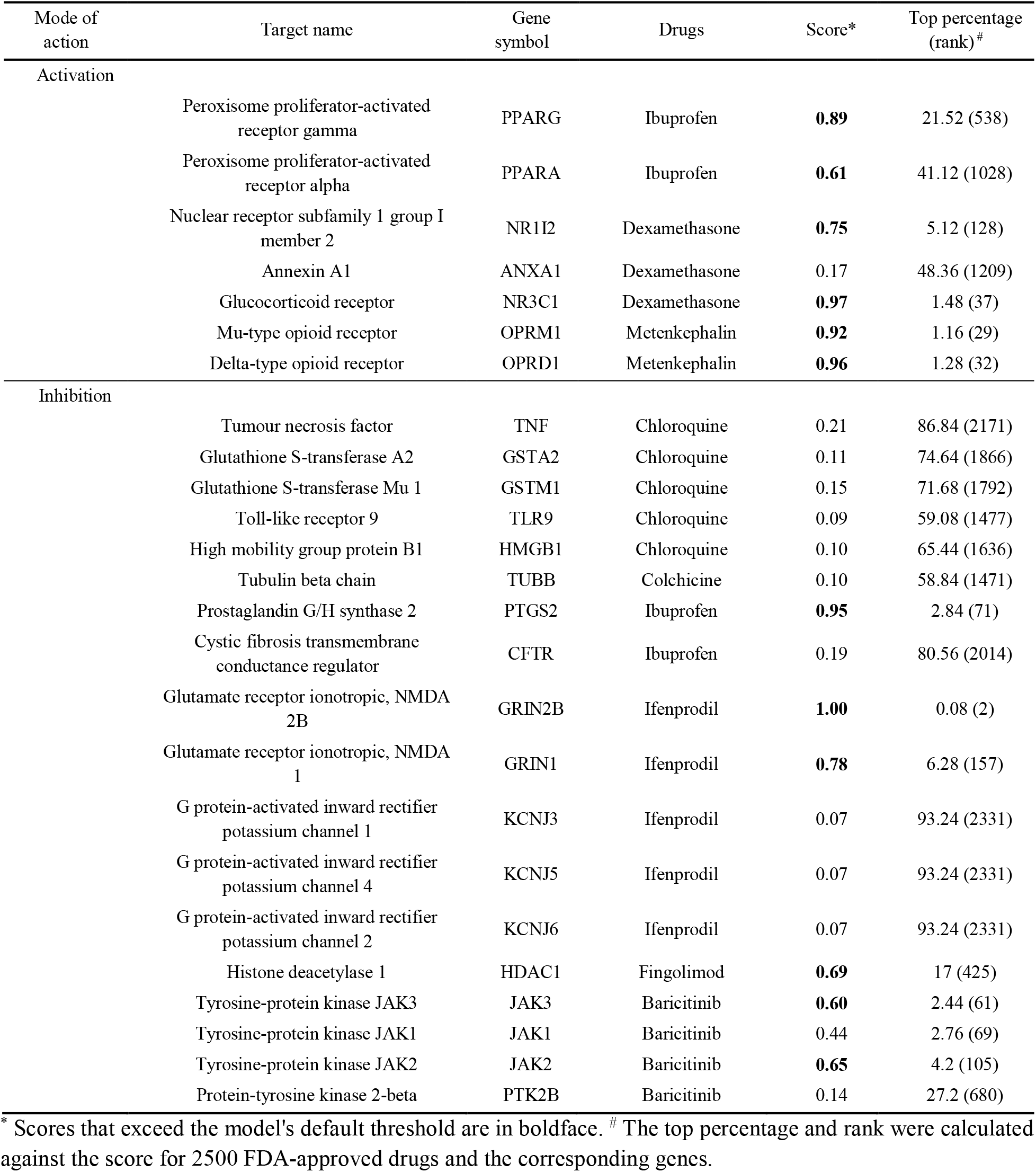
Predicted results for validated DTIs related to COVID-19.

| Mode of action | Target name | Gene symbol | Drugs | Score* | Top percentage (rank)<sup>#</sup> |
| --- | --- | --- | --- | --- | --- |
| Activation | Peroxisome proliferator-activated receptor gamma | PPARG | Ibuprofen | **0.89** | 21.52 (538) |
|  | Peroxisome proliferator-activated receptor alpha | PPARA | Ibuprofen | **0.61** | 41.12 (1028) |
|  | Nuclear receptor subfamily 1 group I member 2 | NR1I2 | Dexamethasone | **0.75** | 5.12 (128) |
|  | Annexin A1 | ANXA1 | Dexamethasone | 0.17 | 48.36 (1209) |
|  | Glucocorticoid receptor | NR3C1 | Dexamethasone | **0.97** | 1.48 (37) |
|  | Mu-type opioid receptor | OPRM1 | Metenkephalin | **0.92** | 1.16 (29) |
|  | Delta-type opioid receptor | OPRD1 | Metenkephalin | **0.96** | 1.28 (32) |
| Inhibition | Tumour necrosis factor | TNF | Chloroquine | 0.21 | 86.84 (2171) |
|  | Glutathione S-transferase A2 | GSTA2 | Chloroquine | 0.11 | 74.64 (1866) |
|  | Glutathione S-transferase Mu 1 | GSTM1 | Chloroquine | 0.15 | 71.68 (1792) |
|  | Toll-like receptor 9 | TLR9 | Chloroquine | 0.09 | 59.08 (1477) |
|  | High mobility group protein B1 | HMGB1 | Chloroquine | 0.10 | 65.44 (1636) |
|  | Tubulin beta chain | TUBB | Colchicine | 0.10 | 58.84 (1471) |
|  | Prostaglandin G/H synthase 2 | PTGS2 | Ibuprofen | **0.95** | 2.84 (71) |
|  | Cystic fibrosis transmembrane conductance regulator | CFTR | Ibuprofen | 0.19 | 80.56 (2014) |
|  | Glutamate receptor ionotropic, NMDA 2B | GRIN2B | Ifenprodil | **1.00** | 0.08 (2) |
|  | Glutamate receptor ionotropic, NMDA 1 | GRIN1 | Ifenprodil | **0.78** | 6.28 (157) |
|  | G protein-activated inward rectifier potassium channel 1 | KCNJ3 | Ifenprodil | 0.07 | 93.24 (2331) |
|  | G protein-activated inward rectifier potassium channel 4 | KCNJ5 | Ifenprodil | 0.07 | 93.24 (2331) |
|  | G protein-activated inward rectifier potassium channel 2 | KCNJ6 | Ifenprodil | 0.07 | 93.24 (2331) |
|  | Histone deacetylase 1 | HDAC1 | Fingolimod | **0.69** | 17 (425) |
|  | Tyrosine-protein kinase JAK3 | JAK3 | Baricitinib | **0.60** | 2.44 (61) |
|  | Tyrosine-protein kinase JAK1 | JAK1 | Baricitinib | 0.44 | 2.76 (69) |
|  | Tyrosine-protein kinase JAK2 | JAK2 | Baricitinib | **0.65** | 4.2 (105) |
|  | Protein-tyrosine kinase 2-beta | PTK2B | Baricitinib | 0.14 | 27.2 (680) |

\* Scores that exceed the model's default threshold are in boldface. <sup>#</sup> The top percentage and rank were calculated against the score for 2500 FDA-approved drugs and the corresponding genes.

## DISCUSSION

Accurately identifying DTIs with a mode of action is a crucial step in the drug development process and understanding the modes of action of drugs. Here, we present AI-DTI, a novel computational approach for identifying activatory and inhibitory targets for small molecules. By leveraging a mol2vec model and genetically perturbed transcriptome, AI-DTI constructs vector representations of activatory and inhibitory DTIs from the structures of small molecules and the responses of biological systems following genetic perturbations. The comprehensive evaluation demonstrated that AI-DTI accurately predicts activatory and inhibitory DTI pairs, even in datasets containing sparse positive samples, DTI pairs that include compounds and/or targets that were unseen in the training phase, and high-throughput biological assay results. A case study of COVID-19 DTIs shows that AI-DTI can be used to prioritize activatory and inhibitory DTIs for novel diseases.

We believe that AI-DTI can bring significant contributions and advantages in two aspects: drug discovery and research on the mechanisms of drugs. In drug discovery, our method can be applied to discover candidate compounds for diseases involving a variety of targets by providing large-scale predictions between a series of small molecules and a wide range of targets. In research on the mechanisms of drugs, our method can generate plausible hypotheses for understanding the mechanisms of action of novel compounds and natural products by predicting activatory and inhibitory DTIs using only the 2D structure of a compound.

Among the employed classifier models, we found that the CDF model yielded the highest performances in our comprehensive experiments. Unlike the deep learning model, the CDF model automatically determines the complexity of the model in a data-dependent way with relatively few parameters and achieves excellent performance across various domains, including simple DTI prediction (7, 9, 26). It is difficult to compare performance directly due to differences in datasets; however, the our method using the CDF model not only outperformed the previous model that predicts activatory and inhibitory DTIs but also competed with some state-of-the-art models that predict only simple interactions while requiring functional annotations of compounds such as drug-drug interactions, drug-disease relationships, and drug side effects (12, 33, 34).

We showed that AI-DTI is a practical tool that accurately predicts DTIs and their modes of action; however, there are several limitations of this study with potential for further improvement. First, the prediction performance can be further improved by applying advanced algorithms, such as GCN, which have been recently reported to show state-of-the-art performance (14). Since the previous model still requires functional annotation of drugs, such as drug-drug interactions, an interesting future study will be to develop a model that predicts DTIs more accurately, even for novel compounds. Second, we used transcriptome profiles transduced with cDNA and shRNA as target vectors, which could include potential off-target effects. The performance of our model may be improved further by upgrades based on large-scale datasets created using advanced techniques, such as CRISPR. A future direction of our work is to develop a versatile predictive model that accurately predicts DTIs with various modes of action.

## DATA AVAILABILITY

AI-DTI is an open source available in the Bitbucket repository (https://bitbucket.org/NNSM/ai_dti)

## SUPPLEMENTARY DATA

Supplementary Data are available at NAR online.

---

**Supplementary Figure 1.**
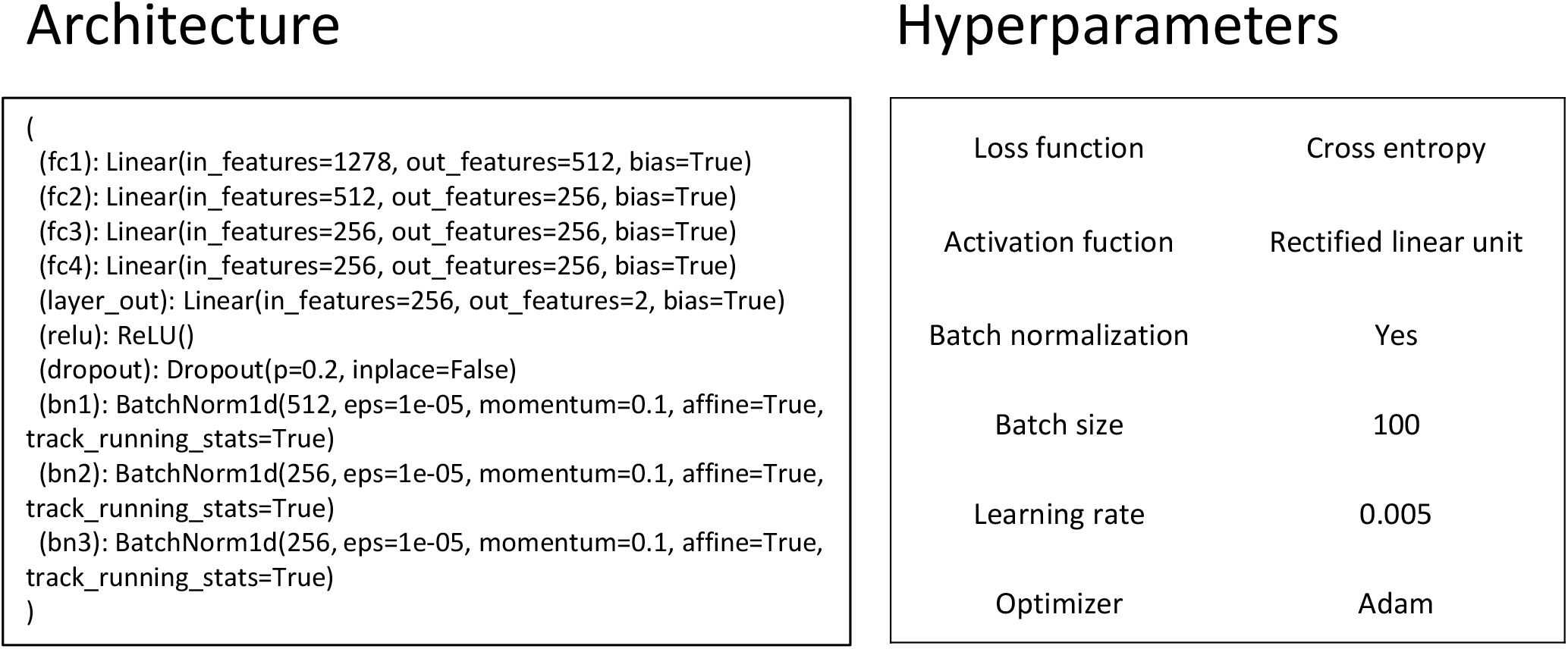
Architecture and hyperparameters of the multilayer perceptron model

## References

1. Hughes, J.P., Rees, S.S., Kalindjian, S.B. and Philpott, K.L. (2011) Principles of early drug discovery. Br. J. Pharmacol., 162, 1239–1249.

2. Kapetanovic, I.M. (2008) Computer-aided drug discovery and development (CADDD): In silico-chemico-biological approach. Chem. Biol. Interact., 10.1016/j.cbi.2006.12.006.

3. Hassan Baig, M., Ahmad, K., Roy, S., Mohammad Ashraf, J., Adil, M., Haris Siddiqui, M., Khan, S., Amjad Kamal, M., Provazník, I. and Choi, I. (2016) Computer Aided Drug Design: Success and Limitations. Curr. Pharm. Des., 10.2174/1381612822666151125000550.

4. Lee, W.Y., Lee, C.Y., Kim, Y.S. and Kim, C.E. (2019) The methodological trends of traditional herbal medicine employing network pharmacology. Biomolecules, 9, 362.

5. Fang, J., Liu, C., Wang, Q., Lin, P. and Cheng, F. (2017) In silico polypharmacology of natural products. Brief. Bioinform., 19, 1153–1171.

6. Mousavian, Z. and Masoudi-Nejad, A. (2014) Drug-target interaction prediction via chemogenomic space: Learning-based methods. Expert Opin. Drug Metab. Toxicol., 10.1517/17425255.2014.950222.

7. Chu, Y., Kaushik, A.C., Wang, X., Wang, W., Zhang, Y., Shan, X., Salahub, D.R., Xiong, Y. and Wei, D.Q. (2021) DTI-CDF: a cascade deep forest model towards the prediction of drug-target interactions based on hybrid features. Brief. Bioinform., 22, 451–462.

8. Zeng, X., Zhu, S., Lu, W., Liu, Z., Huang, J., Zhou, Y., Fang, J., Huang, Y., Guo, H., Li, L., et al. (2020) Target identification among known drugs by deep learning from heterogeneous networks. Chem. Sci., 11, 1775– 1797.

9. Zeng, X., Zhu, S., Hou, Y., Zhang, P., Li, L., Li, J., Huang, L.F., Lewis, S.J., Nussinov, R. and Cheng, F. (2020) Network-based prediction of drug-target interactions using an arbitrary-order proximity embedded deep forest. Bioinformatics, 10.1093/bioinformatics/btaa010.

10. Wang, Z., Zhou, M. and Arnold, C. (2020) Toward heterogeneous information fusion: bipartite graph convolutional networks for in silico drug repurposing. Bioinformatics, 36, i525–i533.

11. Olayan, R.S., Ashoor, H. and Bajic, V.B. (2018) DDR: Efficient computational method to predict drug-Target interactions using graph mining and machine learning approaches. Bioinformatics, 34, 1164–1173.

12. Luo, Y., Zhao, X., Zhou, J., Yang, J., Zhang, Y., Kuang, W., Peng, J., Chen, L. and Zeng, J. (2017) A network integration approach for drug-target interaction prediction and computational drug repositioning from heterogeneous information. Nat. Commun., 8.

13. Lee, I., Keum, J. and Nam, H. (2019) DeepConv-DTI: Prediction of drug-target interactions via deep learning with convolution on protein sequences. PLoS Comput. Biol., 15, 1–21.

14. Zhao, T., Hu, Y., Valsdottir, L.R., Zang, T. and Peng, J. (2020) Identifying drug–target interactions based on graph convolutional network and deep neural network. Brief. Bioinform., 00, 1–10.

15. Zhang, Y.F., Wang, X., Kaushik, A.C., Chu, Y., Shan, X., Zhao, M.Z., Xu, Q. and Wei, D.Q. (2020) SPVec: A Word2vec-Inspired Feature Representation Method for Drug-Target Interaction Prediction. Front. Chem., 7, 1–11.

16. Subramanian, A., Narayan, R., Corsello, S.M., Root, D.E., Wong, B., Golub, T.R., Subramanian, A., Narayan, R., Corsello, S.M., Peck, D.D., et al. (2017) Resource A Next Generation Connectivity Map?: L1000 Platform Resource A Next Generation Connectivity Map?: Cell, 171, 1437–1452.e17.

17. Keenan, A.B., Jenkins, S.L., Jagodnik, K.M., Koplev, S., He, E., Torre, D., Wang, Z., Dohlman, A.B., Silverstein, M.C., Lachmann, A., et al. (2018) The Library of Integrated Network-Based Cellular Signatures NIH Program: System-Level Cataloging of Human Cells Response to Perturbations. Cell Syst., 6, 13–24.

18. Sawada, R., Iwata, M., Tabei, Y., Yamato, H. and Yamanishi, Y. (2018) Predicting inhibitory and activatory drug targets by chemically and genetically perturbed transcriptome signatures. Sci. Rep., 10.1038/s41598-017-18315-9.

19. Jaeger, S., Fulle, S. and Turk, S. (2018) Mol2vec: Unsupervised Machine Learning Approach with Chemical Intuition. J. Chem. Inf. Model., 58, 27–35.

20. Enache, O.M., Lahr, D.L., Natoli, T.E., Litichevskiy, L., Wadden, D., Flynn, C., Gould, J., Asiedu, J.K., Narayan, R. and Subramanian, A. (2019) The GCTx format and cmap{Py, R, M, J} packages: Resources for optimized storage and integrated traversal of annotated dense matrices. Bioinformatics, 35, 1427– 1429.

21. Smith, I., Greenside, P.G., Natoli, T., Lahr, D.L., Wadden, D., Tirosh, I., Narayan, R., Root, D.E., Golub, T.R., Subramanian, A., et al. (2017) Evaluation of RNAi and CRISPR technologies by large-scale gene expression profiling in the Connectivity Map. PLoS Biol., 15, 1–23.

22. Szklarczyk, D., Gable, A.L., Lyon, D., Junge, A., Wyder, S., Huerta-Cepas, J., Simonovic, M., Doncheva, N.T., Morris, J.H., Bork, P., et al. (19) STRING v11: Protein-protein association networks with increased coverage, supporting functional discovery in genome-wide experimental datasets. Nucleic Acids Res., 10.1093/nar/gky1131.

23. Wang, N., Zhao, G., Zhang, Y., Wang, X., Zhao, L., Xu, P. and Shou, D. (2017) A Network Pharmacology Approach to Determine the Active Components and Potential Targets of Curculigo Orchioides in the Treatment of Osteoporosis. Med. Sci. Monit.

24. Wishart, D.S., Feunang, Y.D., Guo, A.C., Lo, E.J., Marcu, A., Grant, J.R., Sajed, T., Johnson, D., Li, C., Sayeeda, Z., et al. (2018) DrugBank 5.0: A major update to the DrugBank database for 2018. Nucleic Acids Res., 10.1093/nar/gkx1037.

25. Zhou, Z.H. and Feng, J. (2017) Deep forest: Towards an alternative to deep neural networks. In IJCAI International Joint Conference on Artificial Intelligence.

26. Zhou, Z.H. and Feng, J. (2019) Deep forest. Natl. Sci. Rev., 6, 74–86.

27. van Laarhoven, T., Nabuurs, S.B. and Marchiori, E. (2011) Gaussian interaction profile kernels for predicting drug-target interaction. Bioinformatics, 10.1093/bioinformatics/btr500.

28. Davis, J. and Goadrich, M. (2006) The relationship between precision-recall and ROC curves. In ACM International Conference Proceeding Series.

29. Huang, C.T., Hsieh, C.H., Chung, Y.H., Oyang, Y.J., Huang, H.C. and Juan, H.F. (2019) Perturbational Gene-Expression Signatures for Combinatorial Drug Discovery. iScience, 10.1016/j.isci.2019.04.039.

30. Noh, H., Shoemaker, J.E. and Gunawan, R. (2018) Network perturbation analysis of gene transcriptional profiles reveals protein targets and mechanism of action of drugs and influenza A viral infection. Nucleic Acids Res., 10.1093/nar/gkx1314.

31. Spreafico, R., Soriaga, L.B., Grosse, J., Virgin, H.W. and Telenti, A. (2020) Advances in genomics for drug development. Genes (Basel)., 10.3390/genes11080942.

32. Rognan, D. (2020) LIT-PCBA: An Unbiased Data Set for Machine Learning and Virtual Screening. 10.1021/acs.jcim.0c00155.

33. Zheng, X., Ding, H., Mamitsuka, H. and Zhu, S. (2013) Collaborative matrix factorization with multiple similarities for predicting drug-Target interactions. In Proceedings of the ACM SIGKDD International Conference on Knowledge Discovery and Data Mining.

34. Zong, N., Kim, H., Ngo, V. and Harismendy, O. (2017) Deep mining heterogeneous networks of biomedical linked data to predict novel drug-target associations. Bioinformatics, 10.1093/bioinformatics/btx160.

